# AlphaImpute2: Fast and accurate pedigree and population based imputation for hundreds of thousands of individuals in livestock populations

**DOI:** 10.1101/2020.09.16.299677

**Authors:** Andrew Whalen, John M Hickey

## Abstract

In this paper we present a new imputation algorithm, AlphaImpute2, which performs fast and accurate pedigree and population based imputation for livestock populations of hundreds of thousands of individuals. Genetic imputation is a tool used in genetics to decrease the cost of genotyping a population, by genotyping a small number of individuals at high-density and the remaining individuals at low-density. Shared haplotype segments between the high-density and low-density individuals can then be used to fill in the missing genotypes of the low-density individuals. As the size of genetics datasets have grown, the computational cost of performing imputation has increased, particularly in agricultural breeding programs where there might be hundreds of thousands of genotyped individuals. To address this issue, we present a new imputation algorithm, AlphaImpute2, which performs population imputation by using a particle based approximation to the Li and Stephens which exploits the Positional Burrows Wheeler Transform, and performs pedigree imputation using an approximate version of multi-locus iterative peeling. We tested AlphaImpute2 on four simulated datasets designed to mimic the pedigrees found in a real pig breeding program. We compared AlphaImpute2 to AlphaImpute, AlphaPeel, findhap version 4, and Beagle 5.1. We found that AlphaImpute2 had the highest accuracy, with an accuracy of 0.993 for low-density individuals on the pedigree with 107,000 individuals, compared to an accuracy of 0.942 for Beagle 5.1, 0.940 for AlphaImpute, and 0.801 for findhap. AlphaImpute2 was also the fastest software tested, with a runtime of 105 minutes a pedigree of 107,000 individuals and 5,000 markers was 105 minutes, compared to 190 minutes for Beagle 5.1, 395 minutes for findhap, and 7,859 minutes AlphaImpute. We believe that AlphaImpute2 will enable fast and accurate large scale imputation for agricultural populations as they scale to hundreds of thousands or millions of genotyped individuals.

## Introduction

In this paper we present a new imputation algorithm, AlphaImpute2, which performs fast and accurate pedigree and population based imputation for livestock populations of hundreds of thousands of individuals. Genetic imputation is a commonly used tool in agricultural and human genetics. It can be used to decrease the cost of genotyping individuals by allowing only a small number of individuals to be genotyped on a high-cost high-density genotyping platform, and the remaining individuals to be genotyped on a lower-cost lower-density platform. Shared haplotype segments between the low-density and the high-density individuals are then used to fill in missing genotypes for the low-density individuals [1,2]. Low cost genotypes are important for increasing the rate of genetic gain in animal and plant breeding programs [3–5]. As genotyping animals has become a routine part of breeding operations, many agricultural datasets contain hundreds of thousands, or even millions of genotyped individuals [6,7] which means that imputation algorithms must be to scale to ever expanding datasets.

Genetic imputation algorithms use either (1) pedigree or family information to perform imputation, which rely on long shared haplotype segments between an individual and their parents, (2) population information to perform imputation, which rely on haplotype sharing between an individual and distant relatives, or (3) both sources of information in a combined algorithm. Pedigree based imputation tends to be fast and accurate, but requires the pedigree of the population to be known, and many of the founders to be genotyped at high density [8–11]. Population based imputation tend to be slower and less accurate, particularly at low marker densities, but can perform imputation on individuals with unknown parents and no known genotyped relatives [2,12]. Population and pedigree based imputation can be effectively combined for livestock populations: pedigree information is used to impute the genotypes of most individuals, and population information is used to impute the remaining genotypes, particularly those of founders or individuals with ungenotyped parents [9,13,14]. When aiming to improve the scaling of a combined imputation algorithm, most of the runtime tends to occur in the population imputation steps [13].

There have been a large number improvements in the runtime of population based imputation algorithms, particularly those based on the “Li and Stephens” hidden Markov model framework [2]. In this framework, an individual’s genotypes are modelled as a mosaic of pairs of haplotypes from the reference library. The reference library represents possible ancestral haplotypes in the population, and generally consists of all of the phased haplotypes of the high-density individuals. By itself, this algorithm scales poorly, with a runtime that is quadratic with the number of haplotypes in the reference library. Runtime can be improved by either using a fixed subset of haplotypes from the reference library [15,16], or by using a phasing algorithm to pre-phase the data and running haploid hidden Markov model separately on each phased chromosome [17,18]. Because the haploid hidden Markov model only needs to consider one chromosome at a time, it scales linearly with the number of haplotypes in the reference panel, allowing it to scale to reference panels of tens of thousands of haplotypes.

For reference haplotypes with hundreds of thousands of haplotypes, scaling can be improved by employing the Positional Burrows Wheeler Transform (PBWT; [19]). The PBWT is an opportunistic data structure which lexicographically sorts the haplotypes at each loci. By sorting the library in this way, it is possible to search through the haplotype reference library for a given haplotype segment in constant time (independent of the size of the library). The creation of the PBWT is linear in both the number of markers and number of individuals, but once created, it can be re-used for all of the individuals genotyped with the same set of markers. There are a growing number of approaches for using the PBWT to speed up the runtime of imputation, e.g., by using it to find a fixed-number of reference haplotypes to use for haploid imputation [20], find “maximally matching” haplotype segments [19], or implement a Viterbi algorithm by using a branch and bound search [21].

In this paper we first present a new population imputation algorithm which uses the PBWT to perform a guided stochastic search through the haplotype reference library. The idea behind this algorithm is to focus on combinations of haplotypes that have high posterior probability. We do this by creating a series of particles and having them explore the high probability paths through the haplotype reference library. Normally the number of particles we would need to use would scale based on the size of the haplotype reference library. We solve this issue by having the particles represent all of the haplotypes in a region with the same genotype state. We then use the PBWT to update each of these particles in constant time, which allows this approach to scale to large reference haplotype libraries.

We also present a refined version of multi-locus iterative peeling which has greatly reduced runtime and memory requirements compared to previous versions [22]. Multi-locus iterative peeling is a probabilistic method for performing pedigree based imputation, that has high accuracy particularly in the presence of genotyping errors [9,22,23]. However, multi-locus iterative peeling has traditionally been too computationally intensive to use for routine imputation, and most pedigree based imputation algorithms use heuristic methods to perform population imputation [8,11]. We found that it was possible to greatly increase the speed of multi-locus iterative peeling by approximating the joint genotype probabilities of an individual’s parents, and by calling the segregation and genotype states when estimating an offspring’s contribution to their parent’s genotypes. These approximations appear to have limited impact on imputation accuracy.

Finally, we present a combined algorithm which integrates the population and pedigree imputation algorithm.

We have implemented the population, pedigree, and combined imputation algorithms in a new software package, AlphaImpute2. We compared the performance of AlphaImpute2 to AlphaImpute [8], AlphaPeel [22], findhap version 4 [24], Beagle 4.1 [25], and Beagle 5.1 [26] on a series of simulated datasets designed to mimic four real pig pedigrees. We find that AlphaImpute2 has high accuracy and low runtimes, achieving an average accuracy of .99 for low-density individuals across all four pedigrees. The runtime for imputing a single chromosome of a pedigree of 107,000 individuals with 5,000 markers was just over two hours. Compared to the other software, AlphaImpute2 had higher accuracy and lower run-times in most situations.

## Materials and Methods

### Population imputation using particles

AlphaImpute2 performs population based imputation using an approximate version of the Li and Stephens algorithm [2]. In the Li and Stephens algorithm models an individual’s genotype is constructed as a mosaic of haplotypes from a haplotype reference panel. This can be implemented in a hidden Markov model where the state space consists of pairs of haplotype identifiers from the reference panel at each loci, and inference is done to find a high-likelihood path, or sequence of haplotype identifiers, through this space. The path can then be used to impute and phase missing genotypes by looking at that state of each haplotype along the path. Because the state space grows quadratically with the size of the reference panel, approximations are needed to perform inference.

Lunter [21] published an algorithm to produce an exact maximum likelihood (Viterbi) path for a diploid Li and Stephens algorithm in constant time. This approaches uses the Positional Burrows Wheeler Transform [19] as an opportunistic data structure that allows searching across many similar haplotypes at the same time. The approach we use here is loosely based on the framework of Lunter [21]. Instead of taking the maximum likelihood path, we instead generate samples from an approximate posterior distribution over all possible paths through the haplotype reference library. The logic behind this choice is that there may be many paths with similarly high likelihoods, and by combining information from multiple samples it may be possible to obtain more accurate genotypes than from any single sample.

To run the algorithm we construct a series of series of a particles. Each particle consists of a pair of ranges of haplotypes at each loci, 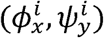 where 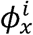 and 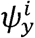 give the set of haplotypes at loci *i*, whose values at preceding loci are given by the sequences *x* = {*x*_*i*−*n*_, *x*_*i*−*n*+1_, …, *x*_*i*_} or *y* = {*y*_*i*−*n*_, *y*_*i*−*n*+1_, …, *y*_*i*_}. 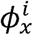 gives the haplotypes for the paternal chromosome, and 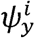 gives the haplotypes for the maternal chromosome. At each loci we probabilistically update the each particle to a new state 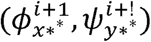 using a guided stochastic search algorithm.

To create the final imputed genotypes and haplotypes, we use this approach to generate between 40-100 particles, and then merge the particles to form a set of consensus haplotypes. These haplotypes can be used to either phase high-density individuals, or impute low-density individuals.

#### Updating a particle

At each loci, particles are updated by probabilistically selecting a new genotype state, and recombination state. We consider a 4×4 grid of possible steps that the particle can take. In the first dimension the particle can move to one of the four phased genotype states. In the second dimension the particle can make the move by having a recombination on either none of their haplotypes, on either their maternal, or paternal haplotypes, or on both haplotypes. An example of this update is given in Figure 1.

**Figure 1:**
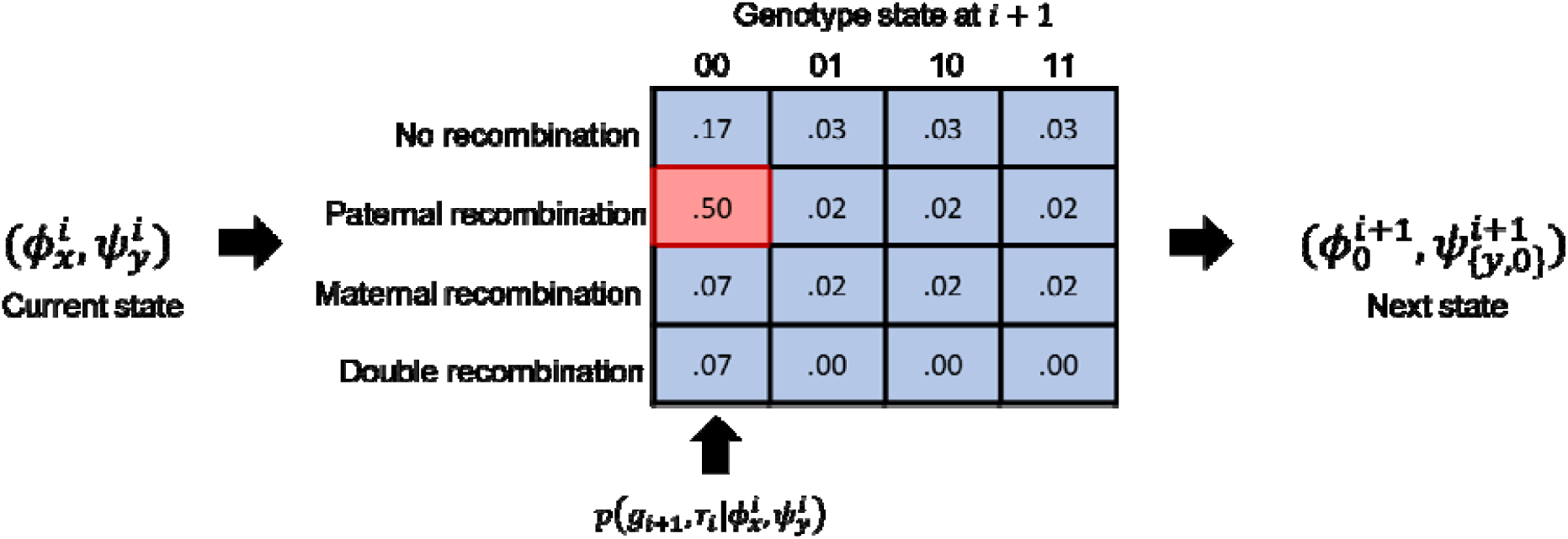
A pictorial representation of a particle update. For each particle at locus we consider a 4×4 grid of possible updates depending on the genotype state at the next locus, and the recombination state between loci. The probability of selecting of each option is given by Equation 1. In the example, the selected state, in red, is homozygous for the reference allele (00), and has a recombination on the paternal haplotype. This means that the paternal path is reset to, while the maternal path is extended to.

The probability of selecting to move to genotype *g*_*i*+1_ with recombination *r*_i_ is:

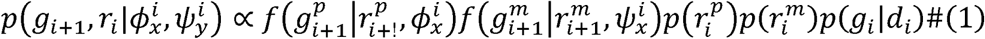

Where 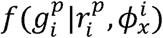 is a function which gives the probability of the paternal part of the genotype state given the recombination state (for either the paternal or maternal haplotypes) and the current set of haplotypes considered; 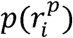 is the probability of the paternal part of the recombination state; and *p* (*g*_*i*_|*d*_*i*_) is the probability of the full genotype state conditional on the data observed; *p* (*g*_*i*_|*d*_*i*_) is the same as in multi-locus iterative peeling and is given later, in Equation 8.

We calculate 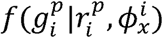 as

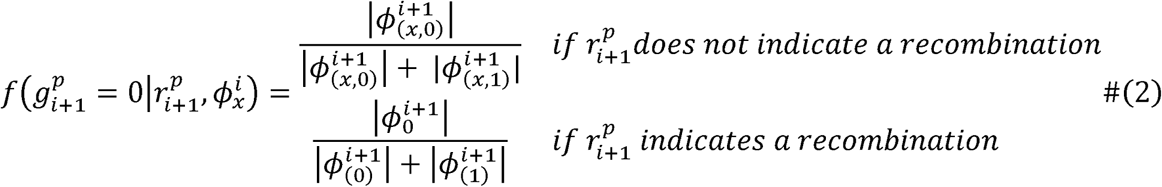

where 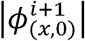 gives the number of haplotypes in 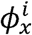 that have a 0 at locus *i* + 1. Similarly 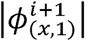 gives the number of haplotypes in 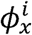 that have a 1 at locus *i* + 1. 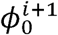 represents the set of all haplotypes that have a 0 at locus *i* + 1. Using the Positional Burrows Wheeler Transform we can calculate 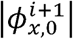 in constant time, independent of the size of the haplotype reference library or the number of haplotypes in 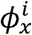 [21].

We calculate the probability of the maternal and paternal recombination 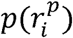 as either:

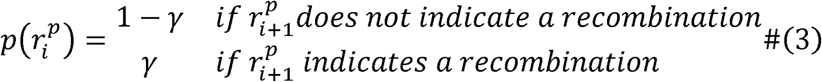

Where γ is a recombination rate which, we estimate as 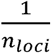. This value underestimates of the effective recombination rate since individuals are inheriting haplotypes from distant relatives [16], however the accuracy of this algorithm seems to largely insensitive to the recombination rate given, and this serves as a good approximation.

In order to determine how well a particle matches the data at a particular locus, we assign a score to each particle at each locus:

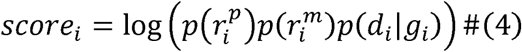

which gives a higher score to particles that do not have a recombination and fit the observed genotype data. We use the score to combine particles into a single consensus haplotype.

#### Combining particles

We create a consensus haplotype and genotype state out of multiple particles by using a two-stage approach. In the first stage we call consensus genotypes for each locus. In the second stage we phase the loci that are called as heterozygous.

To create a consensus genotype, we score each particle at each locus by taking the sum of the scores of each particle (given by Equation 4) within a 50 marker window. We then select the genotype state of the particle with the best score as the called genotype. If multiple particles have the highest score, we use the most frequent genotype state of those particles the called genotype.

In the second stage we phase heterozygous loci by looking at transitions between neighbouring heterozygous states. We track whether the alternative alleles are on the same or different haplotypes, e.g., whether the phased genotype state transitions from *aA* to *aA* where the alternative allele is on the maternal haplotype, or from *aA* to *Aa* where the alternative allele transitions between the maternal to the paternal haplotypes. We pick the transition that is most frequent in all of the particles that are heterozygous at both loci.

#### Backward information

In a traditional hidden Markov model, inference is done by combining information from a forward pass (information from the first locus to the current locus) with a backward pass (information from the last locus to the current locus). The search algorithm we present only takes into account information in the forward pass, and does not make selections based on genotype data from loci after the current locus. Information from backward pass can be useful for imputation by filling in spontaneous missing markers, and phasing genotype states. To incorporate backwards information, we first run a series of particles in reverse, i.e., from the end of the chromosome to the beginning of the chromosome, and then run a forward pass of particles where we replace *p* (*g*_*i*_|*d*_*i*_)in Equation 1 with:

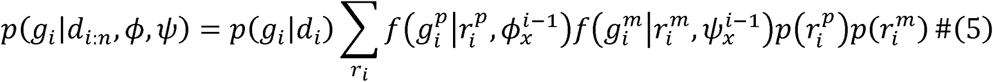

In this equation we project each of the reverse particles forward by one locus to see what genotype state they are likely to carry at the next locus. When multiple particles are run on the backward pass, *p* (*g*_*i*_|*d*_*i:n*_,ϕ,ψ)is averaged across all particles in the backward pass.

#### Imputation

This algorithm can also be used to impute missing markers. To perform imputation, we evaluate particles on the non-missing markers. We track the loci where recombination occurs, and what the set of haplotypes are for each particle at those loci. This creates an ordered pair of haplotype regions {(*start, stop*), 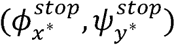}. For each locus, we select the interval that contains the locus, and fill in missing markers on the corresponding haplotypes from the middle haplotype in 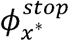 on the paternal side, and 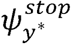 on the maternal side.

#### Creating the haplotype library

In many animal populations pre-phased haplotype libraries are not available and so the haplotype used for population imputation neds to be constructed. This is done by iteratively (1) constructing a haplotype library from the high-density individuals, (2) phasing those individuals with that haplotype library, and (3) re-building the library using the phased haplotypes from the previous iteration. To initialize the haplotype library, we randomly phase heterozygous loci and fill in spontaneous missing genotypes. We then run 5 rounds of phasing and imputation, rebuilding the haplotype library at the end of each round. Running more rounds of phasing and imputation can increase the quality of the haplotype library, but we found that 5 rounds was sufficient in pilot simulations for accurate imputation.

During this process we keep track of location of which haplotypes the individual contributes to the haplotype library, and remove those haplotypes as options for the particle steps. This is done by modifying 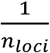 to be either

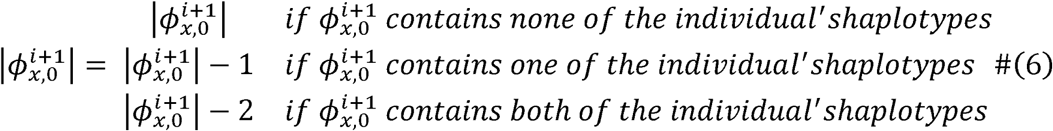

#### Array Clustering

In order to run this algorithm, we want *p* (*g*_*i*_|*d*_*i*_) to be as informative as possible for each locus. Having an informative value for *p* (*g*_*i*_|*d*_*i*_) allows us to have more confidence in taking each step. If the individual does not have genotype data at a locus, then *p* (*g*_*i*_|*d*_*i*_) will be relatively uninformative at that locus. One solution would be to only evaluate the individual’s genotypes at non-missing loci. This would require re-calculating the Positional Burrows Wheeler Transform on an individual-by-individual basis to take into account each individual’s pattern of missing genotypes, which would be prohibitively computationally expensive. Instead we cluster individuals based on the SNP array they are genotyped on, and build the Positional Burrows Wheeler Transform on an array-by-array basis. This greatly reduces the number of times the Positional Burrows Wheeler Transform needs to be calculated, and backwards information is used to guide decisions on any remaining spontaneous missing markers.

### Pedigree based imputation using approximate multi-locus iterative peeling

AlphaImpute2 performs pedigree based imputation using an approximate version of multi-locus iterative peeling [9,22]. In an iterative peeling framework the probability of an individual’s genotypes is based on three sources of information [23]:

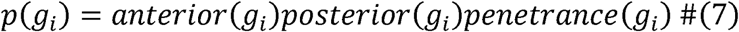

where the anterior term represents information about an individual’s genotypes based on information from the ancestors of the individual filtered through their parents, the posterior represents information about an individual’s genotypes based on the decedents of the individual filtered through their offspring, and the penetrance term represents information about an individual’s genotypes based on their own genetic data (i.e., SNP array or sequence data).

In order to take linkage information into account, multi-locus iterative peeling builds on the normal iterative peeling framework by having the anterior terms and the posterior terms depend on the segregation state of the individual, i.e., which pair of parental haplotypes that individual inherited at each loci [22].

Performing exact inference in a peeling framework is challenging because the anterior and posterior terms for an individual, depend on the genotypes of their parents and offspring, which themselves need to be estimated. Because of this, AlphaImpute2 takes an iterative approach to update the anterior and posterior terms in a series of passes up and down the pedigree. This is summarized in the following algorithm:

1. Downward pass: Starting from individuals in the first generation to the last generation
  a. Re-estimate the segregation probabilities for each individual.
  b. Re-estimate the anterior terms for each individual based on their parent’s genotypes.
2. Upward pass: starting from the last generation to the first generation
  a. Re-estimate the segregation probabilities for each individual.
  b. Re-estimate the posterior terms for each individual based on their offspring’s genotypes.

In order to enable these passes, we sort the pedigree according to an individual’s generation. The generation of each individual is the minimum of the generation of their sire and dam plus one.

The peeling algorithm needs to be run in a series of passes. We have found that 5 passes of peeling is often enough to obtain high-quality genotype probabilities. The purpose of running multiple passes is primarily to transmit information horizontally across the pedigree, i.e., between children of a shared parent. The amount of information that is passed horizontal quickly decays as individuals become more genetically distant.

In order to perform peeling we need to specify how we calculate the penetrance term, and how we update the anterior, posterior, and segregation probabilities.

#### Calculating the penetrance term

The penetrance term give the probability of the observed genetic data, conditional on the individual’s genotype state. We consider four phased genotype states, aa, aA, Aa, AA, where the first allele is the paternal allele, and the second allele is the maternal allele. We assume that the observed genotype data is the number of observed alternative alleles for SNP data.

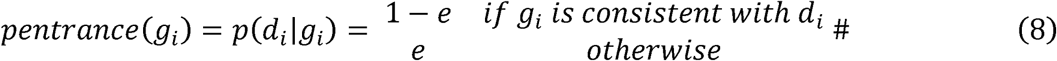

Where *e* represents the genotyping error rate with a default value of 0.01.

#### Updating the anterior term

In each downward pass the anterior term is updated for each individual. To perform this update we re-estimate which genotypes the individual inherited from their parents using the current estimate of their parents’ genotypes. We calculate the anterior term for individual *o* at locus *i* as:

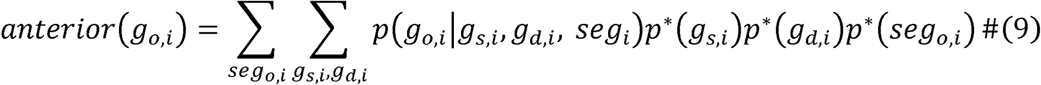

where *p* (*g*_*o,i*_| *g*_*s,i*_,*g*_*d,i*_, *seg*_*i*_) is a transmission function which gives the probability that the offspring inherits a genotype conditional on the offspring’s segregation states and the genotypes of their sire and dam. This value will be either 0 or 1 depending on if an inherited genotype is consistent with the individual’s segregation state and the genotypes of their parents. The term *p* *(*g*_*s,i*_) gives the probability of the genotype state of the sire, based on the information from the last pass of peeling.

This formulation of the anterior term is an approximation to the term used in previous papers [22,23]. In the traditional peeling framework the parents genotypes are given by *p* _−*o*_(*g*_*s,i*_,*g*_*d,i*_) which gives the joint genotype probabilities of the parents ignoring information from the offspring, *o*. By not excluding the offspring’s genotypes we are effectively double counting the offspring’s genotypes: first information on the offspring’s genotypes is used in the penetrance function, and second information from the offspring’s genotypes will be used to calculate the posterior term of the parent, which will then be used to estimate the anterior term of the offspring. In practice, we find that the posterior term from a single offspring only provides a small amount of information to their parent, and that the double counting of information here does not lead to a substantial loss in accuracy. We also assume that the genotype probabilities of the parents are independent, i.e., *p* (*g*_*s,i*_,. *g*_*d,i*_) = *p* (*g*_*s,i*_) *p* (*g*_*d,i*_).

#### Updating the posterior term

In each upward pass the posterior term is updated based on the genotypes of the offspring. This update is performed on a family-by-family basis and the result is combined across families. For a sire, *s*, with mates, *M*= {*m*_1_,*m*_2_,…} the posterior term is:

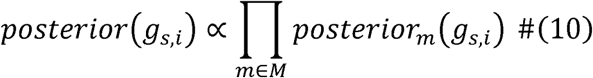

where posterior term for each mate is given by

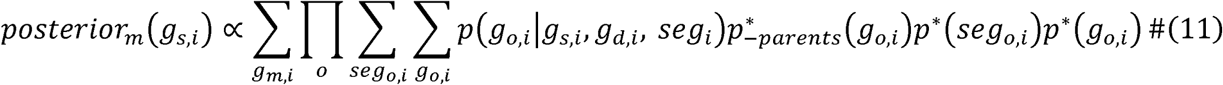

Where the product is on all of the shared offspring between *m* and *s*. Similar terms are used to generate the posterior estimate for each dam.

In order to avoid double counting the genotypes of the parents, we exclude the anterior term from the calculation of the genotypes of the offspring in *p*_−*parents*_ (*g*_*o,i*_). As when calculating the anterior term, we do not exclude the contribution of the offspring when calculating the genotypes of the mate *p* (*g*_*m,i*_).

#### Calculating the probability of each segregation state

The probability of each segregation state are calculated by using a hidden Markov model to determine which segregation state the individual carries at different loci across the genome. We consider four segregation states (*mm, pm, mp, pp*) where the first letter gives whether the individual inherits their sire’s maternal or paternal haplotype, and the second letter gives whether the individual inherits their dam’s maternal or paternal haplotype.

Hidden Markov models are defined by a series of emission probabilities which give the likelihood of the observations given a hidden state, and transmission probabilities, which give the probability of transitioning between hidden states. The emission probabilities of this model are:

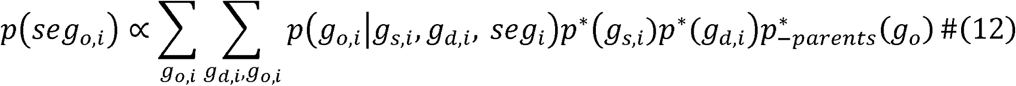

Which uses Bayes’ rule to estimate the probability of each segregation state conditional on the estimated genotypes of the individual and their parents. The transmission function is given by (Whalen 2018):

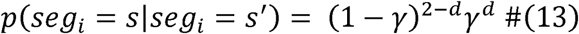

where *d* is the number of recombinations required to move between *s* and *s*′, i.e., *p*(*seg*_*i,j*_ = *pp*|*seg*_*i,j*−1_ = *pm*) = (1 − γ), and γ is the recombination rate. We found that accuracy was largely insensitive to the recombination rate and so set it a default value of 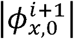 where *n*_*loci*_ is the number of loci on the chromosome. This assumes markers are evenly spaced (in genetic map distance) across an 100 cM chromosome. The assumed total chromosome genetic map length can be changed using a command line option.

We use the forward-backward algorithm [27] to calculate segregation probabilities across each loci. To simplify the amount of information stored at each loci, we assume the segregation probabilities for the maternal and paternal haplotypes are independent and set e.g., *p*(*seg*_*o,pat,i*_ = *m*) = *p*(*seg*_*o,i*_ = *mp*) + *p*(*seg*_*o,i*_ = *mm*).

#### Calling genotype probabilities for the posterior term

In order to reduce runtime we call the offspring segregation and genotype probability values when calculating the posterior terms for their parents. We use calling threshold of 0.99 for calling the segregation values, and a calling threshold of 0.99 for the genotypes on the first round of peeling, and a threshold of 0.95 for subsequent rounds of peeling. For genotype or segregation probabilities that do not reach the threshold, the genotypes or segregation values are set with each state being equally likely.

By calling the segregation values and genotype values, we are able to store part of the posterior update,

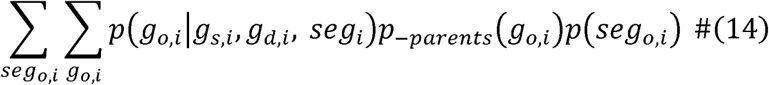

in a look-up table, which substantially reduces runtime. In addition, we do not consider the dependency between the uncertainty of segregation states at nearby loci. The un-modelled linked uncertainty between segregation states can lead to errors in imputing the parental genotypes. By calling the segregation probabilities we mitigate the impact of this simplification by only considering non-equal segregation probabilities where there is minimal uncertainty in the segregation state.

After running the final round of peeling we also call the genotype probabilities of all individuals in the population to get best-guess genotypes for each individual.

### Integrating population and pedigree based imputation

Past work has found that combing pedigree and population imputation algorithms can increase accuracy in populations where pedigree information is available [13,14]. The goal of this combination is to use the population-based imputation algorithms to phase and impute the individuals at the top of the pedigree. These genotypes can then be dropped through the rest of the pedigree using the pedigree based imputation algorithm. To combine the pedigree and population algorithms in AlphaImpute2, we perform imputation using a three step approach where we first perform an initial run of pedigree imputation, we then perform population imputation on a limited set of “pseudo-founders”, and we finish with a final run of pedigree imputation to fill in the remaining missing genotypes.

#### Step 1: Initial pedigree imputation

In Step 1, 5 rounds of multi-locus iterative peeling are run on the population. After the final round all of the genotypes in the population are called with a genotype calling threshold of 0.9. We then split the population to three parts: (1) high-density individuals that have fewer than 10% missing markers, (2) low-density individuals who are “pseudo-founders” (see below) and (3) low-density individuals who are not “pseudo founders”. After splitting the population, the genotypes of individuals in group (3) are reset to their original genotype values before pedigree based imputation.

Pseudo-founders are individuals who genotyped at a higher density than their parents (accounting for the fact that their parents may be imputed to a higher density using pedigree based imputation). To detect pseudo-founders we go through the pedigree from the start to the end and calculate the effective genotyping density of an individual:

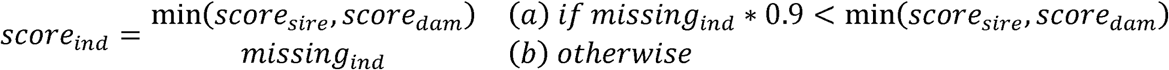

where *missing*_*ind*_ is the percentage of non-missing markers the individual has. The value 0.9 is used to give a slight preference to using the genotype of the parents if the individuals are at a similar marker density. Individuals in group (b) are the “pseudo-founders” of the population.

#### Step 2: Population imputation

In Step 2, we use the population imputation algorithm to phase the high-density individuals detected in Step 1, and use the haplotype constructed from their phased haplotypes as the reference library to impute the low-density “pseudo-founders”. We perform an initial 5 rounds of phasing to iteratively build the reference haplotype library using 40 particles to phase each individual. For imputation we use 100 particles to impute each individual. At the end of this step, we reset the genotypes and haplotypes of high-density individuals that are not “pseudo-founders” to their original genotype states at the start of Step 1. The number of particles selected for phasing and imputation were chosen based on pilot simulations. Larger numbers of particles may yield more accurate results, but the improvement in accuracy will likely be small.

#### Step 3: Final pedigree imputation

In Step 3, we re-run 5 rounds of multi-locus iterative peeling, using the new phased genotypes for the “pseudo-founder” generated in Step 2. In order to reduce the negative impact of switch errors, we perform peeling on a lesioned pedigree where the link between a “pseudo-founder” and both of their parents is removed. After running multi-locus iterative peeling, the genotypes are set to the best-guess genotypes.

### Testing the algorithm

We tested the performance of AlphaImpute2 on four simulated datasets based on pedigrees taken from a commercial pig breeding program. The pedigrees had either 18,349 (*18k*), 34,425 (*34k*), 63,872 (*63k*), or 107,815 (*107k*) individuals and were genotyped on four SNP arrays which ranged from 350 (*very low density*), 10,000 (*low density*), 33000 (*medium density*), and 46,000 (*high density*) markers. Although these marker densities are lower than highest density SNP arrays available for humans and livestock, they represent commonly used marker densities for performing genomic selection in many animal breeding programs [28–30].

We compared the performance of AlphaImpute2 to that of Beagle 4.1, Beagle 5.1, AlphaImpute, AlphaPeel, and findhap. We evaluated each software on their accuracy, runtime, and memory requirements.

### Simulated genetic data

We simulated the four pedigrees by generating a set of founder haplotypes using a Markovian Coalescent Simulator [31], and then dropped them through each pedigrees.

The founder haplotypes were generated by assuming there were 18 100-cM long chromosomes that were simulated using a per site mutation rate of 2.5□×□10−8, and an effective population size (Ne) that changed over time based on estimates for the Holstein cattle population [32]. Ne was set to 100 in the final generation of simulation and to 1256, 4350, and 43,500 at 1000, 10,000, and 100000 generations ago, with linear changes in between. The number of markers per chromosome varied between 1,231 and 4690 based on the marker densities on each chromosome in the real genotype data.

The founder haplotypes were then dropped through the pedigree using AlphaSimR [33]. The genotypes of each individual were then masked to reflect the pattern of missingness for that individual in the real genotype data.

### Comparison with other software

We evaluated the performance of AlphaImpute2 when using either the population only algorithm, the pedigree only algorithm, or the combined algorithm.

We compared the performance of AlphaImpute2 with the performance Beagle 4.1, Beagle 5.1, AlphaPeel, findhap, and AlphaImpute. Beagle 4.1 and Beagle 5.1 were run using default parameters except the effective population size which was set to 200. AlphaImpute and AlphaPeel were run with default parameters. For AlphaImpute we rounded the genotype probabilities that it outputs before calculating accuracy to make it consistent with the other software packages. findhap was run with the recommend parameters of maxlen = 600, minlen = 75, and errate = .004.

Our goal in running a large number of other software packages was to evaluate the performance of both the population only, and pedigree only algorithms separately, and to evaluate the performance of the combined algorithm.

Beagle 4.1 and Beagle 5.1 were chosen to serve as a benchmark for the population only algorithm. Both software packages are commonly used in the human and animal imputation literature, and Beagle 5.1 has incorporated a number of (as of yet unpublished) improvements for phasing.

AlphaPeel was chosen to serve as a benchmark for the pedigree only algorithm. AlphaPeel implements a version of multi-locus iterative peeling, which is approximated by the pedigree only algorithm in AlphaImpute2. Our goal in making this comparison was to see how much accuracy was sacrifice to increase runtime in AlphaImpute2.

findhap and AlphaImpute were chosen to serve as a benchmark for a combined pedigree and population imputation algorithm. Both programs are currently in use in commercial breeding programs, and AlphaImpute2 could serve as a possible candidate to replace them.

### Performance measurements

Imputation accuracy was measured as the correlation between an individual’s imputed genotype and their true genotype, corrected for the population minor allele frequency [34]:

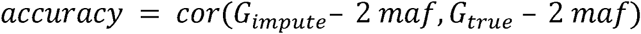

Accuracy was averaged across all of the 18 simulated chromosomes.

We also measured the runtime and memory usage of each program. All programs were run on the Edinburgh Compute and Data Facility cluster using 4 cores. Programs were given at most eight days to impute each chromosome. The results are given only for programs that successfully finished on all of the chromosomes.

## Results

We found that AlphaImpute2 had high accuracy and low run-times across all four pedigrees. Imputation accuracy for the 107k pedigree was .988 for high-density individuals, .988 for medium density individuals, .993 for low-density individuals, and .81 for very-low-density individuals. Imputation took 105 minutes and 14.4 GB of memory for Chromosome 1 (4,600 Markers and 107,000 individuals). AlphaImpute2 had higher accuracy than the alternative algorithms and comparable run times to findhap and Beagle 5.1, both of which are significantly faster than Beagle 4.1. The accuracy, runtime, and memory usage of each algorithm is given in Table 1.

**Table 1:** Imputation accuracy, memory, and runtime for each algorithm. Beagle 4.1 did not complete within 8 days on the 63k or 107k pedigrees. Imputation accuracies were averaged across all 18 chromosomes. Time and memory are given for Chromosome 1.

|  | *High*<br>*Density* | *Medium*<br>*Density* | *Low*<br>*Density* | *Very Low*<br>*Density* | *Time (m)* | *Memory*<br>*(GB)* |
| --- | --- | --- | --- | --- | --- | --- |
| **18k Pedigree** |  |  |  |  |  |  |
| AlphaImpute2 | .998 | .944 | .990 | .827 | 15 | 2.8 |
| *Pedigree Only* | .998 | .661 | .987 | .862 | 3 | 2.5 |
| *Population Only* | .987 | .929 | .963 | .257 | 13 | 2.1 |
| AlphaPeel | .997 | .733 | .984 | .855 | 15 | 9.1 |
| Beagle 4.1 | .995 | .944 | .969 | .327 | 2,250 | 12.4 |
| Beagle 5.1 | .626 | .909 | .939 | .219 | 28 | 17.1 |
| AlphaImpute | .940 | .875 | .931 | .641 | 348 | 10.5 |
| findhap | .719 | .627 | .774 | .445 | 15 | 4.6 |
| **34k Pedigree** |  |  |  |  |  |  |
| AlphaImpute2 | .998 | .961 | .989 | .699 | 24 | 4.2 |
| *Pedigree Only* | .998 | .653 | .987 | .843 | 5 | 4.1 |
| *Population Only* | .992 | .952 | .962 | .199 | 25 | 3.6 |
| AlphaPeel | .997 | .748 | .986 | .838 | 21 | 16.5 |
| Beagle 4.1 | .996 | .959 | .972 | .269 | 4,353 | 17.5 |
| Beagle 5.1 | .601 | .937 | .936 | .148 | 46 | 26.0 |
| AlphaImpute | .955 | .883 | .946 | .611 | 707 | 15.4 |
| findhap | .731 | .678 | .787 | .438 | 36 | 4.6 |
| **63k Pedigree** |  |  |  |  |  |  |
| AlphaImpute2 | .998 | .984 | .991 | .858 | 53 | 8.6 |
| *Pedigree Only* | .976 | .541 | .974 | .864 | 9 | 7.7 |
| *Population Only* | .988 | .978 | .962 | .188 | 57 | 6.3 |
| AlphaPeel | .980 | .596 | .979 | .855 | 74 | 29.8 |
| Beagle 4.1 |  |  |  |  |  |  |
| Beagle 5.1 | .578 | .910 | .937 | .140 | 88 | 26.5 |
| AlphaImpute | .898 | .869 | .921 | .625 | 2,556 | 27.0 |
| findhap | .757 | .732 | .796 | .451 | 156 | 4.8 |
| **107k Pedigree** |  |  |  |  |  |  |
| AlphaImpute2 | .998 | .987 | .993 | .821 | 105 | 14.4 |
| *Pedigree Only* | .991 | .719 | .991 | .869 | 15 | 12.6 |
| *Population Only* | .990 | .981 | .963 | .148 | 106 | 11.6 |
| AlphaPeel | .992 | .786 | .990 | .858 | 84 | 50.4 |
| Beagle 4.1 |  |  |  |  |  |  |
| Beagle 5.1 | .682 | .948 | .942 | .114 | 190 | 43.7 |
| AlphaImpute | .942 | .929 | .940 | .639 | 7,859 | 41.2 |
| findhap | .778 | .761 | .801 | .455 | 395 | 5.0 |

### Accuracy of the full AlphaImpute2 algorithm

The accuracy of AlphaImpute2 depended primarily on the genotyping density of the individuals and their relative position in the pedigree. We found similar accuracies across all four pedigrees and so focus on the 18k pedigree to enable comparisons to Beagle 4.1 which did not finish on all pedigrees.

On the 18k pedigree the accuracy of the full AlphaImpute2 algorithm was .998 for high-density individuals, .944 for medium-density individuals, .990 for low-density individuals, and .827 for very-low-density individuals. The lower accuracy for medium-density individuals compared to low-density individuals was likely driven by their relative position in the pedigree. All of the medium-density individuals appeared in the first quarter of the pedigree compared to only 3% of the low-density individuals and 0.2% of the high-density individuals.

The accuracy of the pedigree only algorithm was .998 for high-density individuals, .661 for medium-density individuals, .987 for the low-density individuals, and .862 for the very-low-density individuals. Compared to the full algorithm, the pedigree only algorithm had much lower accuracy on the medium-density individuals (.661 compared to .944), and similar accuracies on the high-density, low-density, and very-low-density individuals.

The accuracy of the population only algorithm was .987 for the high-density individuals, .929 for the medium-density individuals, .973 for the low-density individuals, and .257 for the very-low-density individuals. Compared to the full algorithm, the population only algorithm had much lower accuracy on the very-low-density individuals (0.257 compared to 0.827), and between 1-2% lower accuracies on the high-density, medium-density, and low-density individuals.

The full AlphaImpute2 algorithm had higher accuracy than both the population only or pedigree only algorithms except in the case of very-low-density individuals where the pedigree only algorithm had a slightly higher accuracy (.862 compared to .827).

### Pedigree only imputation accuracy compared to AlphaPeel

The pedigree only algorithm in AlphaImpute2 uses an approximate version of multi-locus iterative peeling that is implemented in AlphaPeel. The accuracy of the two algorithms are similar, with the accuracy of AlphaPeel on the 18k pedigree being .997 for high-density individuals, .733 for medium-density individuals, .984 for low-density individuals, .855 for very-low-density individuals.

The correlation between the genotypes imputed by the pedigree-only imputation algorithm and AlphaPeel was high. On Chromosome 1 for the 18k pedigree, the correlation between the genotypes imputed between the two algorithms was on average .973, with a correlation of .999 for high-density individuals, .960 for medium-density individuals, .994 for low-density individuals, and .804 for very-low-density individuals. The lower correlation for the medium-density and very-low-density individuals is due to the lack of high-density parents for these individuals. In AlphaPeel, the observed minor allele frequency is used as a prior for the genotypes of the founder individuals, whereas a minor allele frequency of 0.5 is used as a prior for founder individuals in AlphaImpute2. We return to this difference in the Discussion.

### AlphaImpute2 accuracy compared to Beagle 4.1 and Beagle 5.1

For the 18k pedigree, the accuracy of Beagle 4.1 was .995 for the high-density individuals, .944 for the medium-density individuals, .969 for the low-density individuals, and .327 for the very-low-density individuals. The accuracy of Beagle 5.1 was .626 for the high-density individuals, .909 for the medium-density individuals, .939 for the low-density individuals, and .219 for the very-low-density individuals.

The accuracy of Beagle 4.1 was slightly higher than that of Beagle 5.1 in all cases, with the largest difference being on filling spontaneous missing markers in the high-density individuals, where the accuracy of Beagle 4.1 was .995 but the accuracy of Beagle 5.1 was .626.

The accuracy of the population only algorithm in AlphaImpute2 was similar to Beagle 4.1 with lower accuracies on the medium-density individuals (.929 compared to .944), and very-low-density individuals (.257 compared to .327).

### Combined algorithms: findhap and AlphaImpute2

For the 18k pedigree, the accuracy of findhap was .719 for the high-density individuals, .627 for the medium-density individuals, .774 for the low-density individuals, and .445 for the very-low-density individuals. The accuracy of findhap was between 20-40% lower than combined algorithm in AlphaImpute2 in all cases (Table 1).

The accuracy of AlphaImpute2 was .940 for the high-density individuals, .875 for the medium-density individuals, .931 for the low-density individuals, and .641 for the very-low-density individuals. The accuracy of the combined algorithm in AlphaImpute2 was higher than AlphaImpute in all cases, with the largest differences for medium density individuals (.944 compared to .857) and very-low-density individuals (.827 compared with .641).

### Runtime

AlphaImpute2 was faster than all of the other software packages tested. For the pedigree of 18k individuals, AlphaImpute2 took 15 minutes, followed by findhap which took 17 minutes, Beagle 5.1 which took 28.1 minutes, AlphaImpute which took 348 minutes, and Beagle 4.1 which took 2,250 minutes.

For the pedigree of 107k individuals, AlphaImpute2 took 105 minutes, Beagle 5.1 took 190 minutes, findhap took 395 minutes, and AlphaImpute took 7,859. Beagle 4.1 did not finish on the 107k pedigree within eight days of run time.

## Discussion

In this paper we present a new population and pedigree based imputation algorithm, AlphaImpute2, and demonstrate its performance on four simulated datasets based on real livestock pedigrees. We find that it is able to perform fast and accurate imputation in a range of scenarios, and preforms competitively with other existing imputation software. In the remainder of the discussion we discuss the advantages of combining pedigree and population based imputation information for imputation, ways to further decrease the runtime of AlphaImpute2, the performance of the approximate iterative peeling framework, compare the population imputation algorithm to already existing population imputation algorithms, and the particle based approach for approximating the Li and Stephens algorithm.

### Combining pedigree and population imputation increases accuracy

In line with previous research, we find that combining population and pedigree imputation can increase accuracy compared to running either the population or pedigree imputation algorithms alone [13].

Compared to the pedigree only algorithm, the combined algorithm delivers high-accuracy phasing and imputation for the individuals at the top of the pedigree. These phased genotypes can then be used to impute individuals further down in the pedigree. This improves the imputation accuracy for both the “pseudo-founders” of the pedigree, but also other descendants who may be genotyped at lower densities. A similar effect was seen in LDMIP which used a population based imputation algorithm to impute and phase the founders of the pedigree before running multi-locus iterative peeling [9].

Compared to the population only algorithm, the combined algorithm delivers higher-accuracy imputation across the board, particularly for very-low-density individuals. For these individuals imputation accuracy is improved by using pedigree information to decrease the number of haplotypes that need to be considered – the four parental haplotypes in the case of pedigree based imputation, compared to tens of thousands for population imputation – which makes it easier to find the correct haplotypes with a limited number of low-density markers.

The only place where accuracy of the combined algorithm was lower than that of the pedigree only algorithm, was for very-low-density individuals, particularly those at the beginning of the pedigree. The lower accuracy on very-low-density individuals is likely due to a lack of high-density or medium-density ancestors for these individuals. In AlphaPeel, the minor allele frequency is used as a prior for missing genotypes of founders in the population. This allows AlphaPeel to take the uncertainty in the genotypes of these individuals into account. In contrast, in the combined algorithm the founders and “pseudo founders” are imputed with the population imputation algorithm, and the resulting genotypes are treated as observed genotypes (with a default 1% error rate). The population imputation algorithm tends to have low error rates for high, medium, and low-density individuals, but high error rates for very-low-density individuals. Treating these imputed genotypes as observed in the final round of population imputation may be the cause of the lower imputation accuracy. A solution to this problem may be to include a minimum genotyping density required for population imputation (e.g., 10-50 non-missing markers per chromosome), and using the minor allele frequency as a prior for the genotypes of “pseudo founders” who do not reach this density.

### Decreasing the runtime of AlphaImpute2

We found that the combined imputation algorithm had lower runtime than the population only algorithm, but a higher runtime than the pedigree-only algorithm. The lower runtime compared to the population only algorithm is likely due to the fact that in the combined algorithm imputation is only run on a small set of “pseudo founders”. This does not lead to a large reduction in runtime since all the of the high-density individuals are still need to be phased to build the haplotype reference library.

One option to decrease runtime would be to bypass phasing completely by using the high-density individuals who have been fully phased via pedigree imputation to construct the haplotype reference library. We tested this in a small number of pilot simulations and found that this approach reduced run time by 50%, but also decreased accuracy by 1-2%. The lower accuracy is likely driven by having a less relevant set of haplotypes included in the reference library, particularly from those individuals at the top of the pedigree.

Another option to decrease runtime of the population imputation algorithm would be to decrease the number of particles that are run. We chose 40 particles for phasing the haplotype reference panel, and 100 for imputing low-density individuals since those values seemed to give good accuracies in pilot simulations. The number of particles used for phasing the haplotype reference panel was lower than that for imputation, since errors in the haplotype reference panel can be corrected in imputation, and the cost of each additional particle is higher for phasing since phasing is run five times on the high-density individuals to refine the haplotype reference library.

### Approximate iterative peeling

One of the goals in this paper was to improve the scaling of multi-locus iterative peeling. We have previously found that multi-locus iterative peeling is a robust imputation algorithm for performing imputation in large livestock pedigrees [22], but has suffered from long run-times that make it impractical for regular use. We found that by approximating the full multi-locus iterative peeling algorithm we were able to reduce both runtime and memory by 80% by a factor while maintaining the similar accuracies. The speed improvements in AlphaImpute2, exploit the fact that offspring provide relatively little information on their parent’s genotype, and in many cases it is possible to call the segregation values at most loci. This allows us to re-use the parent’s genotype probabilities in the peel-down steps for all of their offspring instead of re-calculating these probabilities on an offspring-by-offspring basis, and to use lookup tables to calculate the summations in the peeling-up step (particularly Equation 11).

In terms of accuracy the AlphaPeel and the pedigree only algorithm in AlphaImpute2 had similar accuracies in both datasets. This suggests that the use of the approximations lead minimal decreases in accuracy on these datasets. The primary difference between algorithms was on how the founders of the population were imputed. For missing genotypes in the founders, AlphaPeel imputes the individuals based on the minor allele frequency in the population. AlphaImpute2 imputes these individuals assuming a minor allele frequency of 0.5. We used a neutral minor allele frequency in AlphaImpute2, to prevent the algorithm from incorrectly calling genotypes with a low minor allele frequency for the combined algorithm, and assume that the genotypes of these individuals will eventually be imputed using the population imputation algorithm. This means that when run alone, the pedigree only algorithm in AlphaImpute2 may give lower accuracies than AlphaPeel, but we find that the combined algorithm in AlphaImpute2 gives higher accuracies than AlphaPeel in most cases.

### Comparison of population imputation algorithms

Compared to the other imputation algorithms tested, AlphaImpute2 obtained generally higher imputation accuracies at lower runtimes on all four simulated datasets.

In terms of speed, we found that AlphaImpute2 and Beagle 5.1 scaled the best out of the software packages tested. findhap had initially low-runtimes on the 18k pedigree, but the performance substantially decreased as the number of reference haplotypes grew larger. The runtime of findhap increased from 15 minutes to 395 minutes between the 18k pedigree and the 107k pedigree, where the runtime of AlphaImpute2 only increased from 15 minutes to 105 minutes. The poorer scaling of findhap is likely due to it searching through the haplotype reference library for each individual, a task that gets harder as more high-density individuals are genotyped. AlphaImpute2 addresses this issue by applying the positional Burrows Wheeler transform to enable constant-time searches through large haplotype reference libraries.

In terms of accuracy, we found that AlphaImpute2 had a higher accuracy than most of the other software tested. For the pedigree only algorithm, AlphaImpute2 had a similar accuracy to AlphaPeel. For the population only algorithm, AlphaImpute2 had a similar accuracy to Beagle 5.1 and Beagle 4.1. For the combined algorithm AlphaImpute2 had a higher accuracy than all of the other algorithms including findhap and AlphaImpute. These results suggest that the approximations used in multilocus peeling had limited impact on imputation accuracy for pedigree based imputation, that the approximate Li and Stephens algorithm used performs as well as other techniques used to fit the Li and Stephens model, and that the way that population and pedigree based imputation are integrated leads to better performance compared to existing software packages.

We were surprised by the large speed improvement between Beagle 4.1 and Beagle 5.1. Beagle 4.1 had the longest runtime of any of the software packages analysed with a runtime of 36 hours on the 18k pedigree, whereas Beagle 5.1 had a runtime similar to AlphaImpute2 with a runtime of just 28 minutes on the 18k pedigree. The improvement in speed between Beagle 4.1 and Beagle 5.1 are impressive, but the changes to the phasing algorithm are (to our knowledge) as yet unpublished. The paper on Beagle 5 [26] only described the improvements to the haploid imputation algorithm which primarily improve speed on imputing whole genome sequence data.

### Particle based approximation to the Li and Stephens model

The particle based implementation of the Li and Stephens model in AlphaImpute2 takes a different approach for increasing the speed of the Li and Stephens algorithm. Previous work has increased the speed of diploid imputation by first pre-phasing the data, and then running a haploid imputation algorithm on the phased haplotypes [18]. The split between phasing and imputation is important, because for many phasing algorithms increased speed by running the algorithm to directly infer the phased genotype (typically groups of heterozygous loci) instead of inferring the underlying haplotypes of origin. This allowed the algorithms to scale better for known loci, but means that that a haploid imputation algorithm needed to be run after pre-phasing the data to fill in missing loci [35,36]. These phasing and imputation algorithms have then been extended to utilize the Positional Burrows Wheeler transform to increase the speed of both the phasing step [37,38] and imputation step [20].

In contrast, the population imputation algorithm in AlphaImpute2 runs phasing and imputation together in a full diploid Li and Stephens model. In order to make this computationally tractable, we approximate the Li and Stephens model using a small number of particles to search for paths with high posterior probability, and use the Positional Burrows Wheeler Transform to update entire sets of paths at once. Our approach is most similar to that of FastLS [21], with the difference that we generate multiple (approximate) samples from the posterior distribution instead of calculating a single maximum-likelihood path. This has the advantage of guaranteeing a constant-time runtime for each particle, and may increase accuracy if multiple paths have similar high posterior probability. We believe that techniques like AlphaImpute2 and FastLS may offer an alternative avenue for performing imputation using a Li and Stephens style model.

## Conclusion

In this paper we present a new imputation algorithm AlphaImpute2, which combines high-accuracy pedigree imputation with high-accuracy population imputation. The pedigree imputation was performed by an approximate form of multi-locus iterative peeling, and the population imputation was performed using a new algorithm which uses particles to approximate a Li and Stephens style hidden Markov models. We find that in four simulated datasets that the algorithm has higher accuracies and lower runtimes compared to other existing imputation software packages, and that it scales well enough to run imputation on hundreds of thousands of pedigree individuals in a matter of hours. We believe that as the size of agricultural populations increase, this software will provided a much needed tool for performing imputation while scaling to the datasets available.

## Declarations

## Ethics approval and consent to participate

Not applicable.

## Consent for publication

Not applicable.

## Availability of data and material

The base dataset used in this study cannot be made available due to commercial considerations. However, a copy of the simulation pipeline using a purely simulated dataset is available from the authors upon a request.

## Competing interests

The authors declare no competing interests.

## Funding

The authors acknowledge the financial support from the BBSRC ISPG to The Roslin Institute BB/J004235/1, from Genus PLC, and from Grant Nos. BB/M009254/1, BB/L020726/1, BB/N004736/1, BB/N004728/1, BB/L020467/1, BB/N006178/1 and Medical Research Council (MRC) Grant No. MR/M000370/1.

## Authors’ contributions

AW and JMH designed the simulation study. AW carried out the simulations and analysed the results. All authors contributed to writing the manuscript.

## Acknowledgements

This work has made use of the resources provided by the Edinburgh Compute and Data Facility (ECDF) (http://www.ecdf.ed.ac.uk).

## Declarations

### Conflicts of interest/Competing interests

On behalf of all authors, the corresponding author states that there is no conflict of interest.

### Ethics approval

Not applicable

### Consent to participate

Not applicable

### Consent for publication

Not applicable

### Availability of data and material

Not applicable

### Code availability

The code used to simulate the data in this study is available from the authors upon a reasonable request. The method, AlphaImpute2, is available from the authors’s website https://alphagenes.roslin.ed.ac.uk/.

## Notes

### Competing Interest Statement

The authors have declared no competing interest.

